# Meta2DB: Curated Shotgun Metagenomic Feature Sets and Metadata for Health State Prediction

**DOI:** 10.1101/2024.10.03.616398

**Authors:** Car Reen Kok, Nisha J. Mulakken, James B. Thissen, Jose Manuel Martí, Ryan Lee, Jacob B. Trainer, Andre R. Goncalves, Hiranmayi Ranganathan, Aram Avila-Herrera, Crystal J. Jaing, Nicholas A. Be

**Affiliations:** Lawrence Livermore National Laboratory, Livermore, CA; University of California, Merced, Merced, CA

**Author notes:** Corresponding author: Nicholas A. Be, Ph.D., Biosciences and Biotechnology Division, Lawrence Livermore National Laboratory, Livermore, CA 94550.

## Abstract

Meta2DB is a curated metagenomic and metadata database that provides structurally consistent microbiome taxonomy feature count tables for 13,897 samples across 84 studies, 23 disease states, and 34 geographical locations. All samples were uniformly processed using a streamlined metagenomic classification pipeline that employs a reference database indexed to contain all sequences across all kingdoms of life that were present in the NCBI Nucleotide (nt) database retrieved on Jan 04, 2023. This pipeline leverages high-performance computing (HPC) resources at Lawrence Livermore National Laboratory and was used to process 50TB of publicly available raw metagenomic sequence data. Extensive metadata curation was carried out through a combination of manual curation and automated parsing, producing a consistent inter-study metadata table specifically structured to facilitate training of ML models for prediction of human health.

## INTRODUCTION

The human microbiome represents a target with tremendous potential for diagnosing and predicting human health. Microbiome features have been indicated as prognostic factors for a range of disease states, including cancer, gut health disruption, neurodegenerative disease, environmental exposure, and mental health. Development of models that leverage microbiome features for prediction of human health states could facilitate tools with clinically actionable utility and identify microbiome-centric variables for therapeutic intervention.

Machine learning (ML), and specifically deep learning (DL), approaches have demonstrated potential in exploring and classifying microbiomes in the context of human health [1, 2]. Convolutional neural networks (CNNs) have demonstrated the capacity to predict various modes of disease outcomes such as disease severity in ulcerative colitis, cirrhosis and inflammatory bowel disease using microbiome-derived data [3-5].

Microbiome data employed for predictive modeling are often derived from metagenomic sequence data generated from the specimens of interest. Such profiling can involve amplicon sequencing, where a specific gene or region is amplified and assigned to microbial taxonomy, most often the bacterial 16S rRNA sequence.

Alternatively, shotgun metagenomic sequencing can be performed, whereby all gDNA within an extracted specimen are sequenced and the resultant sequence reads are aligned to references via a metagenomic classification software platform. The output from such analyses is a feature table, where a taxonomic identifier for a given taxonomic rank (e.g., genus, species) is assigned a numeric value (number of classified sequences), which can then be transformed as desired and used as input for the ML platform of choice.

The high-dimensional nature of microbiome data derived from many features (*p*) alongside small sample sizes (*n*) often result in large *p* and small *n* datasets. This presents a critical challenge in ML modeling, resulting in overfitting and limiting generalizability to new or unseen data. One strategy for sample size expansion is to integrate publicly available datasets across different studies. While health-relevant microbiome datasets are becoming increasingly available, dataset integration for use in a cohesive model training approach is difficult as 1) microbiome feature tables are generated by distinct metagenomic classification platforms that impart systematic biases, 2) independent studies implement distinct quality and sequence filtering steps, and 3) metadata are inconsistently and sparsely represented.

One avenue for addressing this need is to perform a meta-acquisition of the raw sequencing data corresponding to many parallel studies, and to process all such datasets via one consistent metagenomic classification pipeline. The Human Microbiome Compendium [6] and the curatedMetagenomicData data package [7] implements such an approach providing an excellent resource for consistently structured microbiome profiles across multiple studies using 16S rRNA amplicon sequencing and shotgun metagenomic sequencing respectively. The Human Microbiome Compendium pipeline incorporates the DADA2 algorithm [8] to produce Amplicon Sequence Variants for taxonomic identification. The profiles contained in the curatedMetagenomicData data package are produced via MetaPhlAn3 [9], which employs a marker library reference for assigning shotgun metagenomic data, leveraging the benefits of increased taxonomic resolution of shotgun sequence. The marker library facilitates a computationally efficient workflow, though it is limited to the taxonomic entities for which markers are identified and indexed.

Our approach (**Figure 1**) employs a reference database indexed to contain all sequences present in NCBI Nucleotide (nt) from all kingdoms of life [10] (Database available here: https://benlangmead.github.io/aws-indexes/centrifuge). This approach is inherently more comprehensive and high resolution, but computationally intensive. To provide a complementary resource for microbiome-centric model training, we leveraged high-performance computing (HPC) resources at Lawrence Livermore National Laboratory to perform metagenomic classification on 50TB of publicly available raw metagenomic sequence data. We generated structurally consistent microbiome taxonomy feature count tables for 13,897 samples, spanning across 84 studies [11-94]. We further performed extensive metadata curation of all assessed studies through a combination of manual curation and automated parsing, producing a consistent inter-study metadata table specifically structured to facilitate training of ML models for prediction of human health. These curated datasets are available here as supplemental material and will soon be hosted on a website (Meta2DB) whereby a user interface supported by relational databases is available for data exploration.

**Figure 1.**
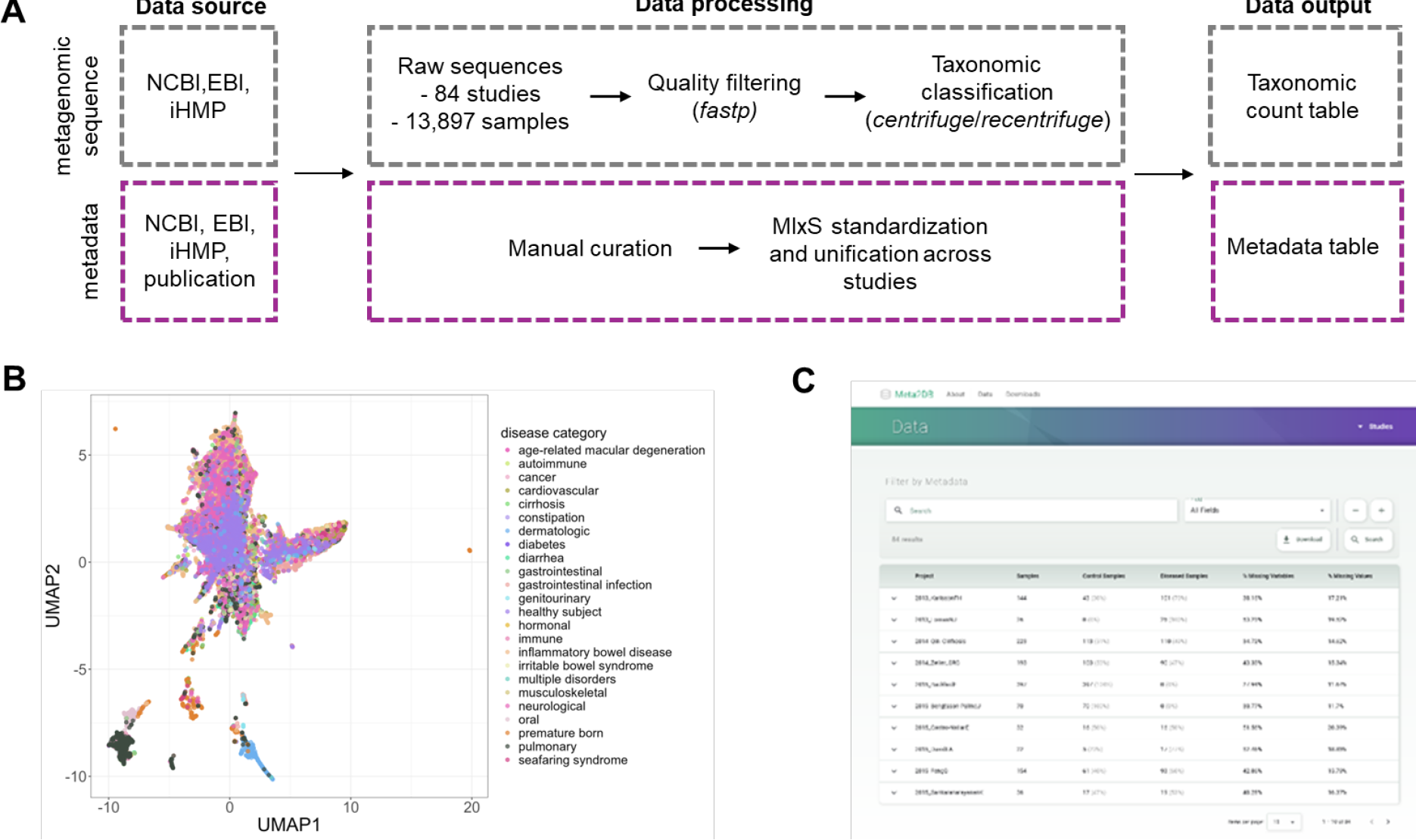
Meta2DB pipeline for the curation of shotgun metagenomic data. A) Data intake, processing and integration of publicly available shotgun metagenomic sequences and associated metadata. B) UMAP plot of 13,897 samples based on computed Euclidean distances. Samples are colored according to different health and disease states. C) Meta2DB web interface for data download and exploration.

## METHODS

### Selection of studies for meta-analysis

Literature studies were selected for data intake and curation if they met each of the following requirements: 1) biospecimens were processed via whole metagenome short-read sequencing using an Illumina platform, 2) raw sequence data and study metadata were publicly accessible, 3) the study was performed in the context of one or more defined human disease states, and 4) each assessed biospecimen could be assigned one of two binary labels corresponding to “control” or “diseased.”

The data collection process includes leveraging an existing resource of curated microbiome data and conducting a strategic search in PubMed. The curatedMetagenomicData package is a pre-existing, well-curated collection of studies, many of which met the characteristics described above [95]. A subset of studies from this curated package was selected for inclusion. To supplement this list of studies, a literature search was carried out in PubMed to retrieve relevant articles. The following search terms were used in combination; “shotgun”, “metagenomics”, “disease”, and studies were manually curated to ensure that they met the requirements listed above.

These parameters were selected for consistency with the aim of facilitating downstream analyses; it is however acknowledged that future analyses could benefit from incorporation of additional data modalities such as amplicon sequences and long-read sequencing, which will be the subject of future iterations.

### Metadata curation

Metadata was extracted from multiple potential sources depending on the individual study of interest, including from the publication and supplementary data or sequence repository. Data sources also included large databases, like the Inflammatory Bowel Disease database of the iHMP, which contains thousands of metagenomics samples from patients with either Crohn’s Disease (CD) or ulcerative colitis (UC), as well as control samples. The Genomic Standards Consortium’s Minimum Information about any Sequence (MIxS) metadata standards were used to group and standardize patient health, treatment, and demographic metadata across all studies. Controlled vocabulary for defined disorders and health states were used to create consistent formatting and naming. In addition to metadata common to all studies, metadata specific to each disorder was also preserved. Metadata around sample collection and type of assay performed were tracked to account for possible effects on outcomes. To distinguish different types of missing data, “not applicable” was used for fields not relevant to a sample, “not available” was used when the data is relevant and missing, and “none” was used when was used when a field that is a treatment is not applied to the patient or when a health status is not observed for the patient.

### Raw sequence acquisition

Raw sequence data (fastq files) were downloaded from public repositories including the NCBI Sequence Read Archive (SRA) [96] and the European Nucleotide Archive (ENA) [97]. The Integrative Human Microbiome Project (iHMP) sequences were obtained from the HMP Data Coordination Center [62].

### Sequence pre-processing

Raw sequence data were pre-processed via fastp [98]. These parameters include using a qualified quality phred value of Q15, an unqualified base percentage of 40%, and a minimum read length of 15. Individual post-filtering of samples was performed with the following read count thresholds: 5M paired-end reads for fecal samples and 1M paired-end reads for skin, oral, and nasal samples. Samples not meeting these thresholds were removed from downstream analysis. Reads were subsequently aligned to the GRCh38 human reference genome using minimap2 [99] for the removal of human reads.

### Metagenomic classification

Resultant reads were processed for metagenomic classification via Centrifuge [100], applying taxonomic labels via Lowest Common Ancestor (LCA) classification strategy. A Minimum Hit Length (MHL) of 15 or 25 was employed for initial Centrifuge processing. Post-processing was performed via Recentrifuge [101] to propagate and normalize Centrifuge assignment scores and perform additional filtering. For purposes of downstream analysis, taxonomic assignments were filtered at a MHL threshold of 40 and feature count tables were produced.

### UMAP visualization

UMAP dimensionality reduction was applied to all 13,897 samples using the umap (version 0.2.10) package in R (version 4.2.2) with default parameters (n_neighbors=15, n_components=2, n_epochs=200). The local neighborhood structure was constructed using the UMAP algorithm on pairwise Euclidean distances.

## Supporting information

Metadata files

Microbiome profiles part 1

Microbiome profiles part 2

Microbiome profiles part 3

Microbiome profiles part 4

Microbiome profiles part 5

Microbiome profiles part 6

Microbiome profiles part 7

Microbiome profiles part 8

Microbiome profiles part 9

## DATA AVAILABILITY

The processed feature tables and metadata are available here as supplemental material. The Meta2DB website will be available soon and will host all processed feature tables, metadata tables along with example notebooks and scripts for data intake and analyses. The indexed version of the centrifuge database based on the entirety of NCBI nt is available at https://benlangmead.github.io/aws-indexes/centrifuge for processing of datasets not included in Meta2DB.

## ACKNOWLEDGEMENTS

This work was performed under the auspices of the U.S. Department of Energy by Lawrence Livermore National Laboratory under Contract DE-AC52-07NA27344. LLNL Disclaimer: This document was prepared as an account of work sponsored by an agency of the United States government. Neither the United States government nor Lawrence Livermore National Security, LLC, nor any of their employees makes any warranty, expressed or implied, or assumes any legal liability or responsibility for the accuracy, completeness, or usefulness of any information, apparatus, product, or process disclosed, or represents that its use would not infringe privately owned rights. Reference herein to any specific commercial product, process, or service by trade name, trademark, manufacturer, or otherwise does not necessarily constitute or imply its endorsement, recommendation, or favoring by the United States government or Lawrence Livermore National Security, LLC. The views and opinions of authors expressed herein do not necessarily state or reflect those of the United States government or Lawrence Livermore National Security, LLC, and shall not be used for advertising or product endorsement purposes.

