## Supplementary material for "Meta2DB: Curated Shotgun Metagenomic Feature Sets and Metadata for Health State Prediction": Metadata files: meta2db_data_definitions.pdf

1. not available:
    a. Data is entered as "not available" for a field when it can be assumed that this data exists and is relevant to a sample, but we do not have this information.
    b. Select examples:
        i. Sample collection related fields: geo_loc_name, collection_date, env_biome, env_feature
        ii. Host related fields: special_diet, host_disease_stat, host_tot_mass, host_height, host_diet, host_body_mass_index, host_smoker, ihmc_ethnicity

2. not applicable:
    a. Data is entered as "not applicable" for a field when the field is not relevant for the sample
    b. Select examples
        i. When seq_type="whole genome sequencing", the fields target_gene and target_subfragment are entered as "not applicable"
        ii. When host_body_product="feces", the host_body_site field is "not applicable"
        iii. When health_disease_stat="control", many disease related fields will be "not applicable". Some examples: cirrhosis_hbv, inr, pt
3. none:
    a. Data is entered as "none" for a field where we have information that something was not observed or was not given to the host. This is most often used for treatment and testing related fields
    b. Select examples
        i. he field: "none" entered for hosts who were tested for hepatic encephalopathy, but had no observed hepatic encephalopathy
        ii. antiviral_medication: "none" entered for hosts that we know were not given antivirals
        iii. beta_blocker: "none" entered for hosts that we know were not given beta blockers
